# Interactions between age and sex in multiscale entropy and spectral power changes across the lifespan

**DOI:** 10.64898/2026.04.09.717268

**Authors:** Jack P. Solomon, Simon G. J. Dobri, Kelly Shen, Vasily A. Vakorin, Sylvain Moreno, Anthony R. McIntosh

## Abstract

Multiscale entropy (MSE) changes in relation to age, whereby aging is associated with an increasing bias towards fine scale entropy. This change is thought to represent a shift toward localized information processing in the brain as we age. However, this relationship has not been tested in large sample sizes alongside other demographic factors and cognitive behaviours. This study aimed to validate previously reported effects of aging on MSE in a large open access database (Cambridge Centre for Ageing and Neuroscience, N=587) and expand the findings to include an investigation of the effects of sex and a variety of cognitive behaviours. MSE curves and power spectrum densities (PSD) were calculated for each region of interest from the magnetoencephalography data. Multivariate partial least squares analyses were used to assess the relationship between MSE or PSD and 5 behavioural / demographic factors including: age, sex, fluid intelligence, visual short-term memory and a generalized measure of cognitive function. Age was associated with increased fine scale and decreased coarse scale entropy, as well as complementary spectral changes, including slowing of peak alpha rhythms, increased beta-band activity, and reduced gamma-band activity, which replicates prior MSE and PSD findings. In both domains, these age-related patterns differentiated based on sex with advancing age. Importantly, the unique effects of sex diverged between MSE and PSD. This result indicates that entropy-based measures can isolate aspects of temporal organization that are not clearly summarized by spectral structure alone.

## Introduction

Aging alters the organization and dynamics of the human brain. While the structural and functional features of neurodegeneration have been extensively characterized, less is known about how brain dynamics evolve across the healthy lifespan (Coelho et al., 2021; Escrichs et al., 2020; Li et al., 2015; Zhong and Chen, 2022). Understanding these normative changes is essential for distinguishing adaptive from pathological aging and identifying individuals at risk of cognitive decline (Chen et al., 2024; Yang et al., 2021). Brain function can be understood as a complex adaptive system in which local and distributed processes interact across multiple spatial and temporal scales (Costa et al., 2005; Deco et al., 2011; McDonough and Nashiro, 2014). These dynamics are constrained by anatomical connectivity but remain flexible, supporting the integration and segregation of information required for complex cognition (Heuvel et al., 2009; Honey et al., 2009). The analysis temporal structure of electrophysiological neural signals therefore provides a promising avenue into brain function that extends beyond traditional measures of oscillatory power (Garrett et al., 2013; Miskovic et al., 2016; Tognoli and Kelso, 2014).

The richness of information conveyed in the temporal domain can be interpreted from its complexity (Costa et al., 2005). Estimates of signal complexity can be attained by a wide variety of calculations, each with their own strengths and weaknesses, see (Sun et al., 2020) for a review covering widely applied methods of entropy in neuroimaging. Among the available complexity measures, multiscale entropy (MSE) is particularly well suited to capturing these properties. MSE quantifies the regularity of time series across multiple temporal scales, which leverages the relationship between time scale and entropy to highlight that correlated noise is more complex than uncorrelated noise at large time scales (Costa et al., 2005; Courtiol et al., 2016; McIntosh, 2019). As a result, MSE can effectively distinguish uncorrelated noise from biologically meaningful variability in complex systems when contrasting against surrogate data. Fine time scale entropy reflects local and long-range fluctuations, while coarse time scale entropy mainly reflects long-range dynamics. During development, MSE increases, reaching a peak in early adulthood, whereas typical aging is associated with relatively greater fine-scale and reduced coarse-scale entropy, which may be interpreted as a shift from integrated to more segregated network organization (Courtiol et al., 2016; McIntosh et al., 2014, 2008). These trends are reduced in individuals at-risk or with cognitive impairment, underscoring MSE’s potential as a sensitive biomarker of mental state (Bertrand et al., 2016; Ghanbari et al., 2015).

To contextualize MSE as a marker of biological brain aging, it is necessary to establish its relationship with key demographic and cognitive variables. Cognitive performance declines heterogeneously across individuals, influenced by factors such as sex and education (Alty et al., 2023; Buckner et al., 2005; Lindenberger, 2014; Salthouse, 2019). Recent studies suggest that MSE is positively correlated to fluid intelligence and the spatiotemporal distribution of this relationship is moderated by sex (Dreszer et al., 2020; Makani et al., 2022). However, these findings are based on relatively small samples, and it remains unclear how MSE patterns generalize across the adult lifespan or interact with multiple cognitive domains.

The present study uses a large, open-access MEG dataset to investigate how multiscale entropy varies with age, sex, and cognitive performance in healthy adults (Shafto et al., 2014; Taylor et al., 2017). Specifically, the objectives of the study are to: validate previously reported age-related patterns of increasing fine-scale and decreasing coarse-scale entropy; quantify how MSE differs with sex and across behavioural measures related to generalized cognitive function, fluid intelligence and visual short-term memory; and compare MSE results with frequency-domain analyses, given the theoretical link between coarse-graining and low-pass filtering in the power spectrum.

We hypothesized that: i) age would be associated with increased fine-scale and decreased coarse-scale entropy; iia) MSE would differ between male and female participants, though the direction of this difference is uncertain given conflicting reports (Kumral et al., 2020; Zhong and Chen, 2022); and iib) sex would moderate the relationship between fluid intelligence and MSE. Lastly, we expected that the representation of aging effects would be present in both the MSE and PSD data but the representation of these effects would differ between modalities given MSE’s sensitivity to the temporal organization of biological signals.

## 2 Methodology

### 2.1 Participants

The data used in this study were acquired from phase 2 of the Cambridge Centre for Ageing and Neuroscience (CamCAN) dataset (available at http://www.mrc-cbu.cam.ac.uk/datasets/camcan/; Shafto et al., 2014; Taylor et al., 2017). From this dataset, the current study used the 8:40 (minutes:seconds) eyes closed resting-state MEG scan available for 646 participants, which was collected in compliance with the Helsinki Declaration and was approved by the Cambridgeshire 2 Research Ethics Committee (reference: 10/H0308/50; Shafto et al., 2014). Participants were uniformly distributed across age deciles (range 18-88) with 363 female and 283 male participants. We analyzed all participants with MEG data and, to ensure that source localization was methodologically consistent, participants were excluded if their structural MRI data were unavailable. Simon Fraser’s Human Research Ethics Board approved the analysis for this study (REB#: 30000954).

### 2.2 Data acquisition

The MEG data were acquired using a 306-channel VectorView MEG System (Elekta Neuromag, Helsinki). The signal was sampled at 1kHz with an online bandpass filter from 0.03-330Hz and head position was continually monitored using four head position indicator coils (Shafto et al., 2014). Electro-ocular (vertical and horizontal) and electrocardiographic recordings were concurrently taken using sets of bipolar electrodes. Anatomical landmarks (nasion, bilateral preauricular points) and distributed points across the scalp were digitized to facilitate co-registration with the participant’s structural MRIs. The T1-weighted MPRAGE sequence was performed using a 2,250ms repetition time and 2.99ms echo time, with a 9° flip angle, 256X240X192mm field of view, and 1mm^3^ voxel size.

### 2.3 MRI pre-processing

Participants’ cortical surface was reconstructed from their MRI data using FreeSurfer’s *recon-all* algorithm (version 7.4.1; Collins et al., 1994; Dale et al., 1999; Desikan et al., 2006; Fischl et al., 2004, 2002, 2001, 1999b, 1999a; Fischl and Dale, 2000). The output of this process was used to define a boundary element model of the brain to facilitate a subsequent calculation of the forward solution (see: 2.5 Source Localization). Participants’ anatomical landmarks and scalp digitization data were used to align the MEG and MRI data using co-registration provided by (Bardouille et al., 2019), which relies on a semi-automated process (MNE-Python coreg, v.0.14; co-registration data available at https://github.com/tbardouille/camcan_coreg).

### 2.4 MEG pre-processing

For this study, we used MEG data from CamCAN release005 that was pre-processed using the Elekta temporal signal space separation filter and interpolation for rejected MEG channels (Shafto et al., 2014). All analyses were performed using Python version 3.11.5 and MNE Python version 1.6.1 (Gramfort et al., 2013). Head motion correction was not applied to the data nor was the data translated to a common headspace. The resting state data was trimmed to 0:30-6:30 (minutes:seconds), a 1-90Hz bandpass filter was applied with a notch filter at 50Hz to attenuate power line noise. An independent component analysis was then applied to reject artifacts associated with cardiac and ocular events using the Picard method (Ablin et al., 2017). This process cleaned the MEG data, which was downsampled to 500Hz and used for subsequent source localization.

### 2.5 Source Localization

A boundary element model was created using the FreeSurfer watershed algorithm for each participant to improve the accuracy of the source model (Ségonne et al., 2004). This model was used to define a forward model with a grid spacing of 5 mm on the participant’s cortical surface using the coregistrations (Bardouille et al., 2019). The source space activity was estimated using a unit-noise gain minimum variance beamformer (Sekihara and Nagarajan, 2008). The data covariance was calculated from a two-minute window of the cleaned data. The noise covariance was calculated from a two-minute window of the empty room recording for each participant, which was pre-processed in the same manner as the resting state recording. The rank of the data was assumed from the output of Maxwell filtering and the orientation of the forward solution was allowed to freely vary to maximize the filter power with a regularization parameter of 0.05. Timeseries were extracted for 68 regions of interest (ROI; 34 per hemisphere) on the cortical surface defined by the Desikan-Killiany Atlas and represents the first right-singular vector of a singular value decomposition of the time courses within each label. The orientation of the vectors are then aligned using a sign flip if needed (Desikan et al., 2006; Gramfort et al., 2013).

Finally, a randomly selected two-minute section of the source localized time series for each participant was used in the subsequent analyses. This window was broken into 30 non-overlapping 4 second epochs for each ROI. The multiscale entropy, frequency decompositions and phase randomization analyses were performed on a per epoch basis and averaged to return 68 timeseries (one per ROI) for each participant.

### 2.6 Multiscale Entropy

Multiscale entropy is the estimation of sample entropy across a range of increasingly coarse grained timeseries. The MSE calculations were performed on each epoch according to (Costa et al., 2005, 2002) which were translated into python from the algorithm available at www.physionet.org/physiotools/mse/. Briefly, this calculation involves two steps, a coarse-graining procedure and the calculation of sample entropy at each level of time scaling. The coarse-graining procedure repeatedly down-samples the MEG timeseries data from each participant at each ROI. At each level of time scaling, sample entropy is calculated to estimate the rate of information generated a time series using a box-counting method. SampEn identifies how often patterns of m (m=2) successive data will remain within a similarity criterion (defined by a threshold r = 0.5 X S.D.) when compared to the rest of the time series (*p^m^*) and then assessing how many of these patterns remain similar when the next sample m+1 is added to the pattern of successive data point (*p^m+1^*). Sample entropy is defined as ln(*p^m^*/*p^m+1^*) the natural log of the ratio of a pattern of consecutive data points *P^m+1^* remain within the similarity criterion, provided that they were close at *p^m^* (Equation #1).

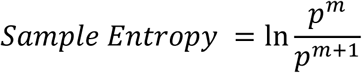

**Equation #1:** Sample entropy as calculation.

One alteration to the original calculation was made for instances where every pattern comparison for p^m^ or p^m+1^ exceeded the threshold r at any given scale. This scenario would result in the sample entropy calculation being undefined (all comparisons for *p^m+1^* exceed the threshold r) or infinite (all comparisons for *p^m^* exceed the threshold r). Costa et al. avoid this scenario by reporting a special case returning the maximal sample entropy for the given coarse-grained vector, see Equation #2a. However, given that the absolute distance between the pattern and other elements in the time series is used, we posit that the maximal sample entropy should be defined by Equation #2b as the distances are sign-invariant. This correction does not affect the calculation of sample entropy outside of this specific scenario.

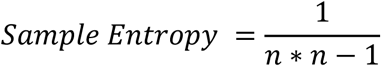

**Equation #2a:** Maximal sample entropy as calculated by Costa et al. 2005 for instances where sample entropy cannot be defined or is infinite.

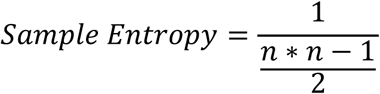

**Equation #2b:** Maximal sample entropy calculation used in this article for instances where sample entropy cannot be defined or is infinite.

### 2.7 Frequency decomposition

In this analysis, we computed spectral power from each ROI using the Welch method with a segment length of 250 using a hamming window. The resulting power spectrum was averaged across epochs and normalized within the ROI as a z-score.

### 2.8 Behavioural Measures

Measures used for our analyses included age, sex, fluid intelligence (Accuracy across four subtests of the Cattell fluid intelligence test; Cattell), visual short-term memory (see (Zhang and Luck, 2008); VTSM) and generalized cognitive function (Addenbrooke’s Cognitive Examination revised; ACE-R) (Cattell, 1987; Mioshi et al., 2006; Zhang and Luck, 2008). Due to the known high correlation between fluid intelligence and age (r=-0.66), scores on the Cattell test were adjusted to represent the residuals from a linear regression model that predicted fluid intelligence from age (Horn and Cattell, 1967). Sex was arbitrarily coded as female = 1 and male = 0. Age was separated into 3 groups based on terciles: Young (18.5-45 years), Middle (45.1-65.6 years) and Old (65.7-88.92 years). The behavioural scores were broken into two groups based on a median split: ACE-R (Low Scoring = 76-97 & High Scoring = 98-100), adjusted Cattell (Low Scoring = -16.16-0.13 & High Scoring = 0.13-12.22 and VSTM (Low Scoring = 0.34-0.79 & High Scoring = 0.80-1.57).

### 2.9 Surrogate analysis with phase randomization

As part of the comparison between MSE and the frequency domain analysis, it was vital to quantify the impact of non-linear dynamics that are uniquely captured by MSE. The impact of non-linear relationships on MSE data, was determined by randomizing the phase of the Fourier transformed signals according to Theiler et al. 1992 and MSE was calculated on the resulting time series (Theiler et al., 1992). The method of generating the phase randomized time series kept the same autocorrelation and power spectrum as the original signal, but the phase dependency was abolished, eliminating nonlinearities from cross-spectral interactions (Courtiol et al., 2016).

### 2.10 Partial Least Squares (PLS) Analysis

We employed two versions of PLS, mean-centered task and behavioural, in this analysis. Mean-centered task PLS was used to investigate overall differences in the MEG data (PSD or MSE) across the demographic factors of interest: age and sex (McIntosh and Lobaugh, 2004). Participants were divided into three age groups: Young [0-33], Middle [34-66], and Old [67-100]. PLS decomposes the group-averaged data using singular value decomposition, which yields mutually orthogonal Latent Variables (LV). An LV is defined as two singular vectors, one containing weights for brain measures (e.g., MSE or PSD) and the other weight for group that index the differences captured by the LV. The strength of the association between the singular vectors is given by its singular value. To investigate potential interactions between the effects of age and sex, separate task PLS analyses were run on each grouping variable (i.e. age alone, sex alone). Latent variables from each of these analyses were compared to the full factorial design by calculating the cosine similarity between the LVs from both models. The same approach was used to assess within-group sex differences for each age tercile.

To test the strength of the relationship between the MSE curves or PSD and our behaviours of interest (fluid intelligence, visual short-term memory, and generalized cognitive function) we applied behavioural PLS analyses to determine patterns of maximal covariance (LV) between the two data matrices. This test allows us to assess the strength of the correlations between each behaviour and the identified patters across the brain. Grouped behavioural PLS analyses were subsequently used to assess group splits based on sex or age would change in the relationships between the MEG data and behaviour matrices in comparison to the ungrouped analysis. The group splits were based on sex, a tercile split of age data and a median split of the three behavioural variables. As was done for the task PLS analysis, the LVs identified in the grouped analyses were compared against the ungrouped test by calculating the cosine-similarity between the LVs from each model. Finally, to assess the similarity of LVs derived from the MSE and power spectrum data, the cosine-similarity of the behavioural LVs between data sets were assessed.

A final set of PLS analyses were used to determine the degree to which MSE effects were driven by linear versus nonlinear dependencies. A behavioural PLS analysis was performed on the MSE curves generated from the phase randomized data, which eliminates systematic nonlinear effects in the data, while preserving linear effects (Theiler et al., 1992). The significant LVs identified in this analysis were compared to the LVs from the identical analysis performed on MSE curves from the unaltered data using their cosine-similarity. If non-linear relationships were dominating differences in the MSE curves, the LVs from both analyses would align poorly. Secondly, a task PLS analysis was used to determine the impact of phase randomization on the MSE to by testing for mean differences in the MSE curves generated from the phase randomized and unaltered data.

Statistical reliability of the PLS results was assessed in two ways. Firstly, null hypothesis testing was done with a permutation test with 10,000 iterations to assess the strength of the singular value for each LV, whereby significance was denoted if the observed singular value falls in the 2.5% tails of the distribution of LVs following permutation (P<0.05; McIntosh and Lobaugh, 2004). This test serves as an “omnibus” test to assesses if the overall relation between the MEG data and behaviours is significant. Secondly, bootstrap estimated standard errors, derived from 10,000 iterations, were used to assess the reliability of the weights within the singular vectors^49^. A ratio of the weight to the standard error (bootstrap ratio) > 3, akin to a 99% confidence interval, was used to identify MSE scale or frequencies that showed the highest reliability. This complements the statistical inference provided by the permutation test by identifying the contribution of individual MEG features to the overall correlation. Finally, scores for each participant were derived from the dot product of the singular vector and the MSE/PSD data. Group differences on an LV were depicted as the average score within group, with the corresponding confidence interval derived from bootstrap estimation.

## 3 Results

Following exclusions (outlined in Methods: Participants) the final sample size was 587 participants. Twenty-two participants were removed for not having MRI data, 8 participants did not have co-registration data, and 18 participants were removed for missing behavioural data. A further 9 participants’ MRI data failed FreeSurfer’s recon_all pipeline and 2 participants’ data failed the MEG pipeline.

### 3.1 Behaviours

After cleaning the data, there were 90-105 participants spread across the 6 demographic groups: young (male = 94, female = 101), middle-aged (male = 95, female 101) and old (male = 105, female = 91). The Cattell scores were adjusted given the high negative correlation with age (see: 2.8 Behavioural Measures). In this sample, the VTSM and ACE-R scores were also negatively correlated with age (Table 1). After adjusting the Cattell scores, the three cognitive behaviours were all positively correlated with each other (Table 1). Two tailed t-tests were performed to test for group differences associated with sex (α=0.05). Age did not differ between the sexes, but female participants had higher VSTM (t=3.91, p=1*10^-4^) and ACE-R (t=2.68, p=0.01) scores and male participants had higher Adjusted Cattell Scores (t=2.21, p=0.03).

**Table 1:** Correlation table (Pearson’s correlation coefficient) between age and behavioural variables (Adjusted Cattell, Visual short-term memory [VTSM] and generalized cognitive function [ACE-R]). Significance is denoted p<0.05*, p<0.01**.

|  | Age | Adjusted Cattell | VTSM | ACE-R |
| --- | --- | --- | --- | --- |
| Age |  | 0.01 | -0.30** | -0.37** |
| Adjusted Cattell | 0.01 |  | 0.25** | 0.45** |
| VTSM | -0.31** | 0.25** |  | 0.37** |
| ACE-R | -0.37** | 0.45** | 0.37** |  |

### 3.2 Validating age related changes in Multiscale Entropy (MSE)

The full factorial mean-centered task PLS with six groups (Age [Young, Middle & Old] X Sex [Male & Female]), revealed two significant LVs that accounted for 98.14% of the covariance between the factorial design and the MSE curves (Figure 1A). The first LV demonstrated an effect of aging, with regional increases in fine-scale MSE and decreases in coarse-scale MSE with age. The second LV highlighted a sex by age effect whereby old male participants have lower fine-scale entropy and increased mid-scale entropy compared to the other groups (Figure 1A).

**Figure 1:**
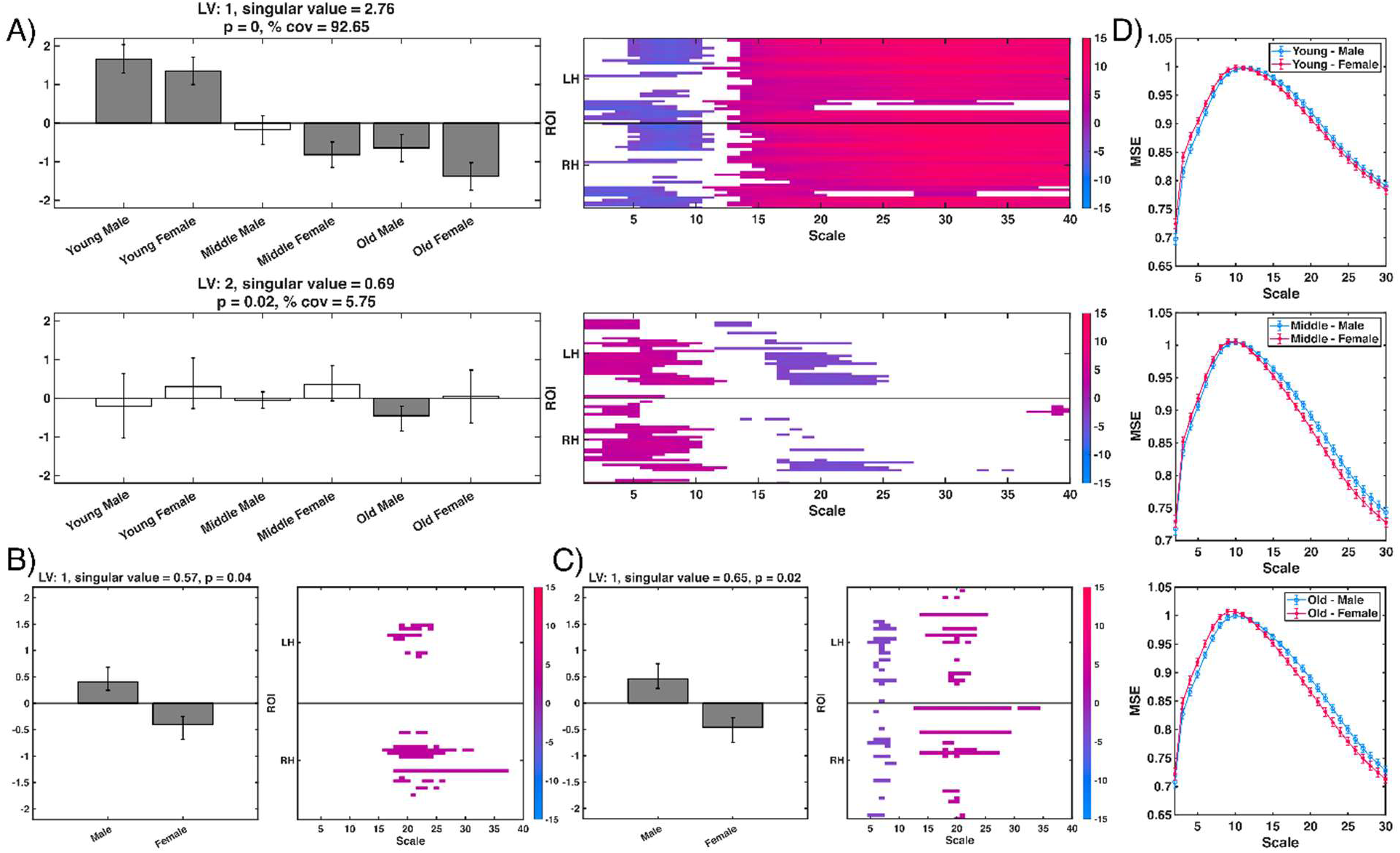
Results of the task partial least squares (PLS) analysis based on the multi-scale (MSE) curves. A) Each significant latent variables (LV), LV1 and LV2, is represented by two plots: 1) a bar plot of the brain scores of each group to the LV ± a 95% confidence interval and 2) a color plot of the bootstrap ratios for the MSE curves from each region of interest (68 total, 34 in each hemisphere; RH = Right hemisphere, LH = Left hemisphere) indicating where the group effects were most reliable. B&C) Results of a task PLS analysis on the middle-aged participants and old participants respectively. The presentation of the significant LVs follows the formatting of subplot A. D) Descriptive plots of the peak of each multiscale entropy curve [mean ± standard error]. Each subplot represents the young, middle aged and old participants (in descending order) and each plot has two series, one for each sex.

The effects of age and sex were further investigated by looking at the between and within group effects of each variable. The between group task PLS analyses revealed a single significant LV for age (singular value = 1.83, p > 1*10^-4^) and sex (singular value=0.49, p=3.3*10^-3^). The age effect aligned highly with LV1 of the full-factorial task PLS (cos(LV1_age_ X LV1 _full factorial_) = 1) whereas the sex effect was captured across both LVs of the full factorial task PLS (cos(LV1_sex_ X LV1 _full factorial_) = 0.71; cos(LV1_sex_ X LV2 _full factorial_) = -0.70). The within-group analyses of age did not reveal a significant LV in the young participants (singular value = 0.46, p = 0.11), but returned the middle and old groups revealed a significant LV highlighting sex differences in each age tercile (Figures 1B & C). These effects were similar to both LVs of the full factorial analysis (cos(LV1_middle_ X LV1 _full factorial_) = 0.80, cos(LV1_middle_ X LV2 _full factorial_) = - 0.51 & cos(LV1_old_ X LV1 _full factorial_) = 0.70, cos(LV1_old_ X LV2 _full factorial_) = -0.66). Male participants showed higher mid-scale entropy in both age groups and female participants had higher fine-scale entropy in the old age group (Figure 1D). These findings would suggest that: 1) the effects of age on MSE validate previous findings and are captured entirely on the first LV of the full factorial design, 2) sex differences are split across both significant LVs, indicating an interaction of age by sex and, 3) sex effects are more noticeable in middle aged and old participants.

Given the established relationship between MSE and the power spectrum, the task PLS analysis was replicated with the power spectrum data. The full factorial PLS with six groups revealed two significant LVs and one marginal LV that accounted for 92.37% of the covariance in the PSD results (Figure 2A). The first LV demonstrated an effect of aging whereby there was cortex-wide alpha slowing (increased activity in the 7-8Hz bin), increased beta (14-30Hz) and low gamma (31-40Hz), along with decreased delta (0-4Hz), theta (5-6Hz) and high gamma bands (40Hz+; Figures 2A&B). The second LV highlighted a sex effect whereby female participants appear to have decreased alpha and high gamma as well as increased beta in comparison to male participants (Figure 2A&C).

**Figure 2:**
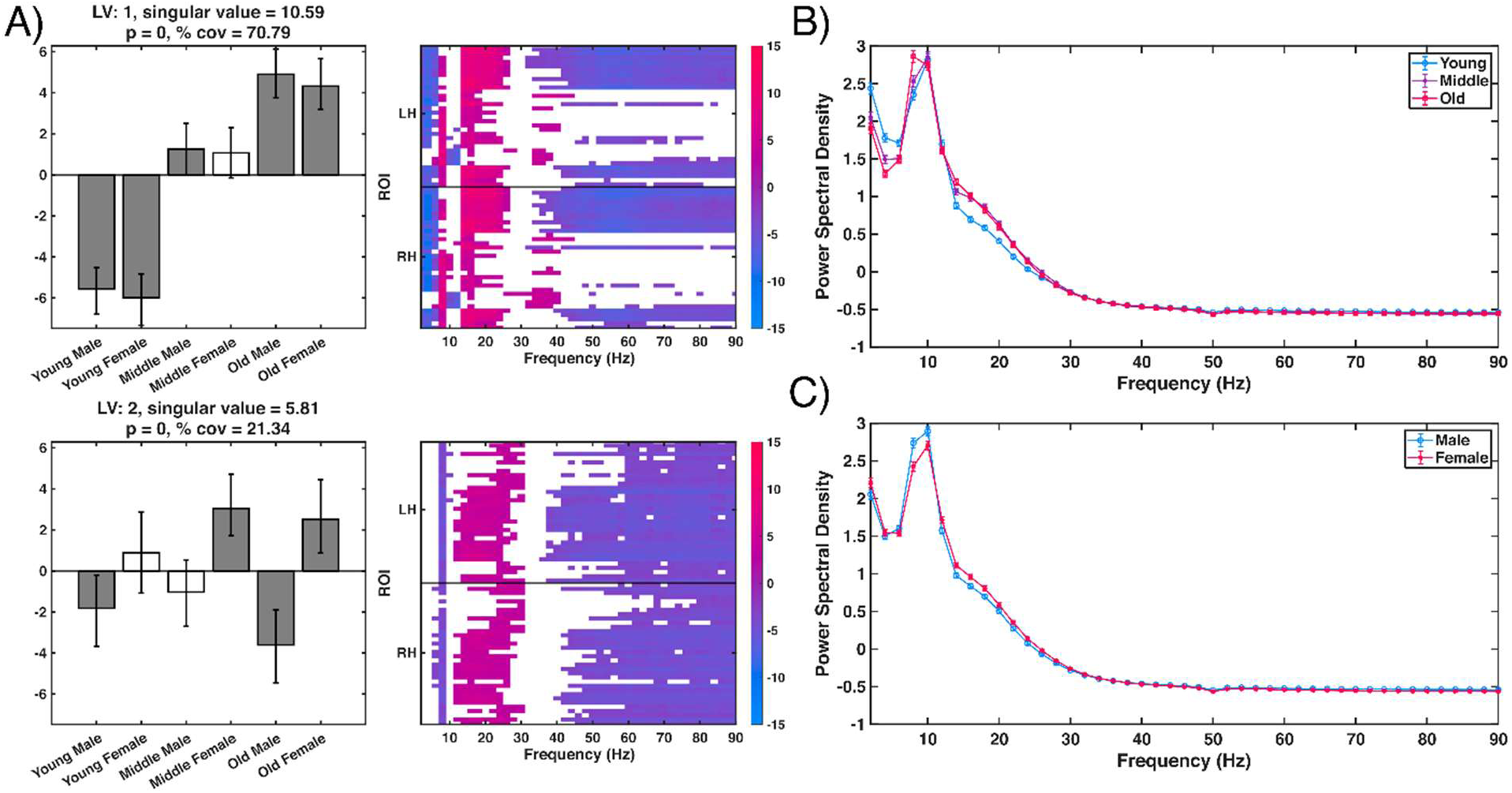
Results of the power spectrum task partial least squares analysis. A) Each significant LV, 1 and 2, are represented by two plots: 1) a bar plot of the brain scores of each group to the LV ± a 95% confidence interval and 2) a color plot of the bootstrap ratios for the power spectrum from each region of interest (68 total, 34 in each hemisphere; RH = Right hemisphere, LH = Left hemisphere). B&C) Descriptive plots of the power spectrum grouped by age and sex respectively [mean ± standard error].

The task PLS analysis of the between group differences in age revealed one significant LV and a marginally significant LV (LV1 singular value = 7.56, p = 0.00; LV2 singular value = 2.12, p = 0.06). The first LV was an aging effect that was highly similar to the first LV of the full factorial design (cos(LV1_age grouped_ X LV1 _full factorial_) = 1), whereas the second LV highlighted a difference in the middle aged participants that was similar to both the second and third LVs of the full factorial analysis (cos(LV2_age grouped_ X LV2 _full factorial_) = 0.62, cos(LV2_age grouped_ X LV3 _full factorial_) = -0.73).

A task PLS of the between group differences associated with sex highlighted a single significant LV representing a difference between male and female participants (singular value = 3.24, p = <1.0X10^-4^) that was most similar to the second LVs of the full factorial design (cos(LV1_sex grouped_ X LV1 _full factorial_) = 0.24, cos(LV1_sex grouped_ X LV2 _full factorial_) = -0.95, cos(LV1_sex grouped_ X LV3 _full factorial_) = -0.17).

Since the sex effects in the PSD results present differently between the age groups, we tested the within group effects of sex in each age tercile using independent task PLS analyses. This analysis identified a significant effect of sex within each age group, with the effect becoming increasingly large with age (Young: singular value = 2.92, p = 0.048; Middle: singular value = 3.15, p = 0.04; Old: singular value = 4.59, p = 0.00). In the young group, the sex effect was most similarly aligned to the second and third LVs in the full factorial PLS (cos(LV1_young_ X LV2 _full factorial_) = -0.66, cos(LV1_young_ X LV3 _full factorial_) = -0.63). In contrast, the sex effect in the two older groups was only similar to the second of the full factorial PLS (cos(LV1_middle_ X LV2 _full factorial_) = -0.92 & cos(LV1_old_ X LV2 _full factorial_) = -0.96). Collectively, these findings suggest that: 1) the effects of age are captured on the first LV of the full factorial design, 2) sex effects are captured by the second (middle aged and old participants) and third (young participants) LVs and 3) sex effects become more robust and widespread with age and reflect differing patterns in the PSD findings.

### 3.3 Relationship between MSE, age, sex and cognitive behaviours

The behavioural PLS analysis investigating the relationship between age, sex, fluid intelligence, visual short-term memory, generalized cognitive function, and the MSE curves returned two significant LVs representing 97.84% of the covariance between behaviour and MSE (Figure 3A). The expression of these effects in the MSE curves for each ROI can be clustered by scale. For the first LV, there were two effects: a cortex-wide relative decrease in coarse-scale entropy and a relative regional increase in fine-scale entropy, largely confined to the frontal and occipital cortices. In the second LV, there was a more widespread relative increase in fine-scale entropy across the parietal and temporal lobes and a relative regional decrease in mid-scale entropy in temporal and occipital lobes.

**Figure 3:**
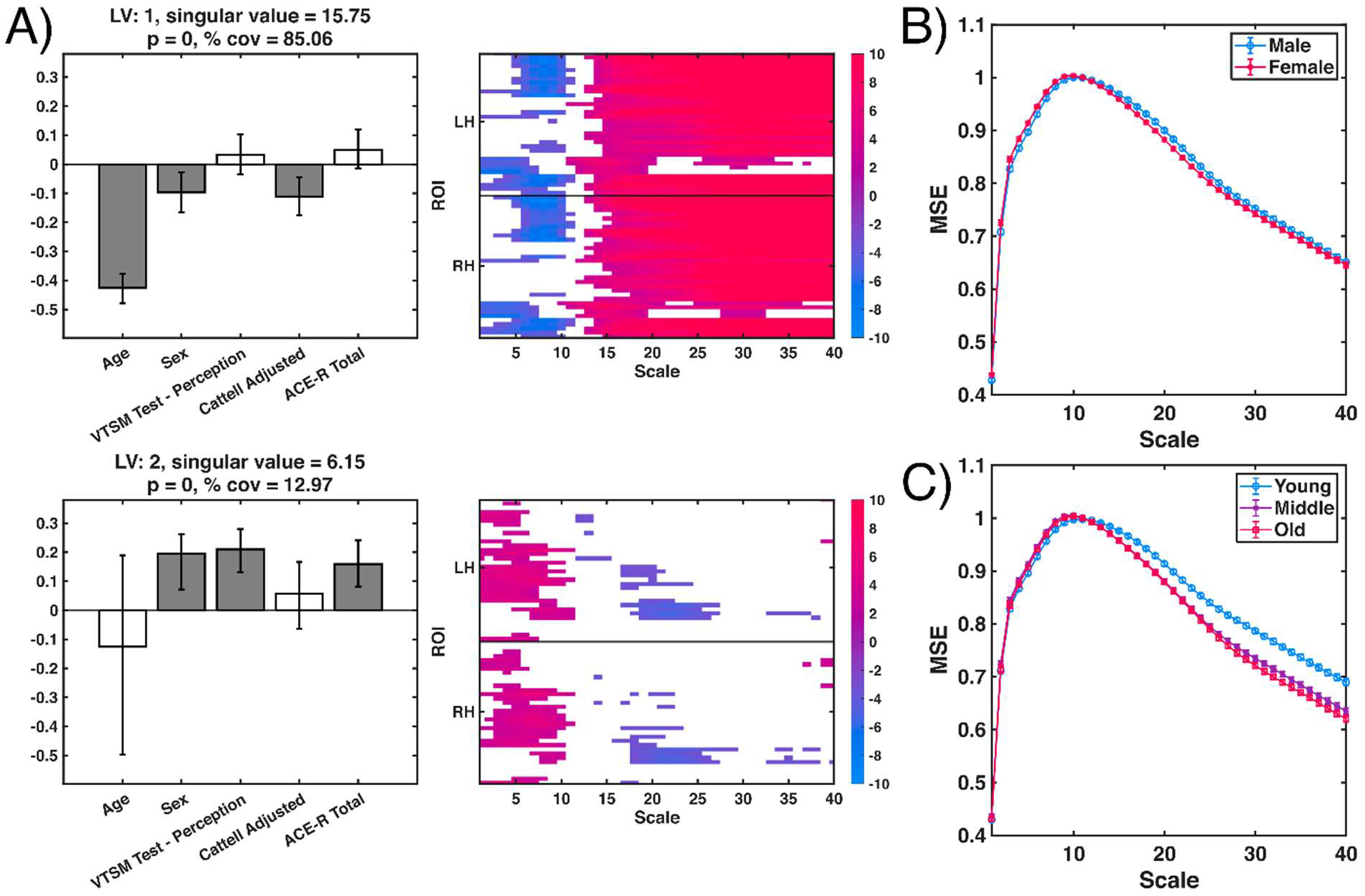
Results of the MSE behavioural partial least squares analysis. A) Each significant LV, 1 and 2, are represented by two plots: 1) a bar plot of the correlation of each behaviour to the LV ± a 95% confidence interval and 2) a color plot of the bootstrap ratios for the MSE curves from each region interest (68 total, 34 in each hemisphere; RH = Right hemisphere, LH = Left hemisphere) indicating where correlations with behaviour, age, and sex were most reliable. B & C) Descriptive plots of multiscale entropy curves [mean ± standard error] grouped by age and sex respectively.

The first LV was reliably correlated with age, which aligns with the task PLS results and replicates previous studies demonstrating increased entropy in the very fine and fine scales and a decrease in coarse-scale MSE (Figure 3B). This LV was also correlated with fluid intelligence and sex; however, sex was weakly related to the effect as the magnitude of the correlation between the first LV and sex was r<0.1 (Figure 3A). The correlation between sex and the second LV indicates that, in the involved regions, female participants express higher fine-scale, lower mid-scale and mixed effects in coarse-scale MSE (Figure 3C). This effect was also correlated with short term memory and generalized cognitive function (Figure 3A).

In the power spectral data, the behavioural PLS analysis revealed two significant LVs accounting for 94.91% of the covariance explained by the model (Figure 4). The first LV was reliably correlated with age, sex and fluid intelligence. This LV showed common effects of aging, including decreased delta and theta power, increased beta power, and decreased gamma power, which were robust across the cortex (top row, Figure 4; Gola et al., 2012). The second LV was negatively correlated with age and positively correlated with sex, visual short-term memory and generalized cognitive function. This effect was related to changes in each canonical frequency band (bottom row, Figure 4). The negative correlation of this LV with age indicated that the decrease in alpha band frequency is the standard slowing of peak alpha seen with age (bottom row, Figure 4; Scally et al., 2018; Trammell et al., 2017). There are two noticeable effects in the beta band: regional decreased power in low beta and increased power in higher beta frequencies across most regions of the cortex (bottom row, Figure 4). Lastly, this LV was associated with regional decreases in gamma frequencies (bottom row, Figure 4).

**Figure 4:**
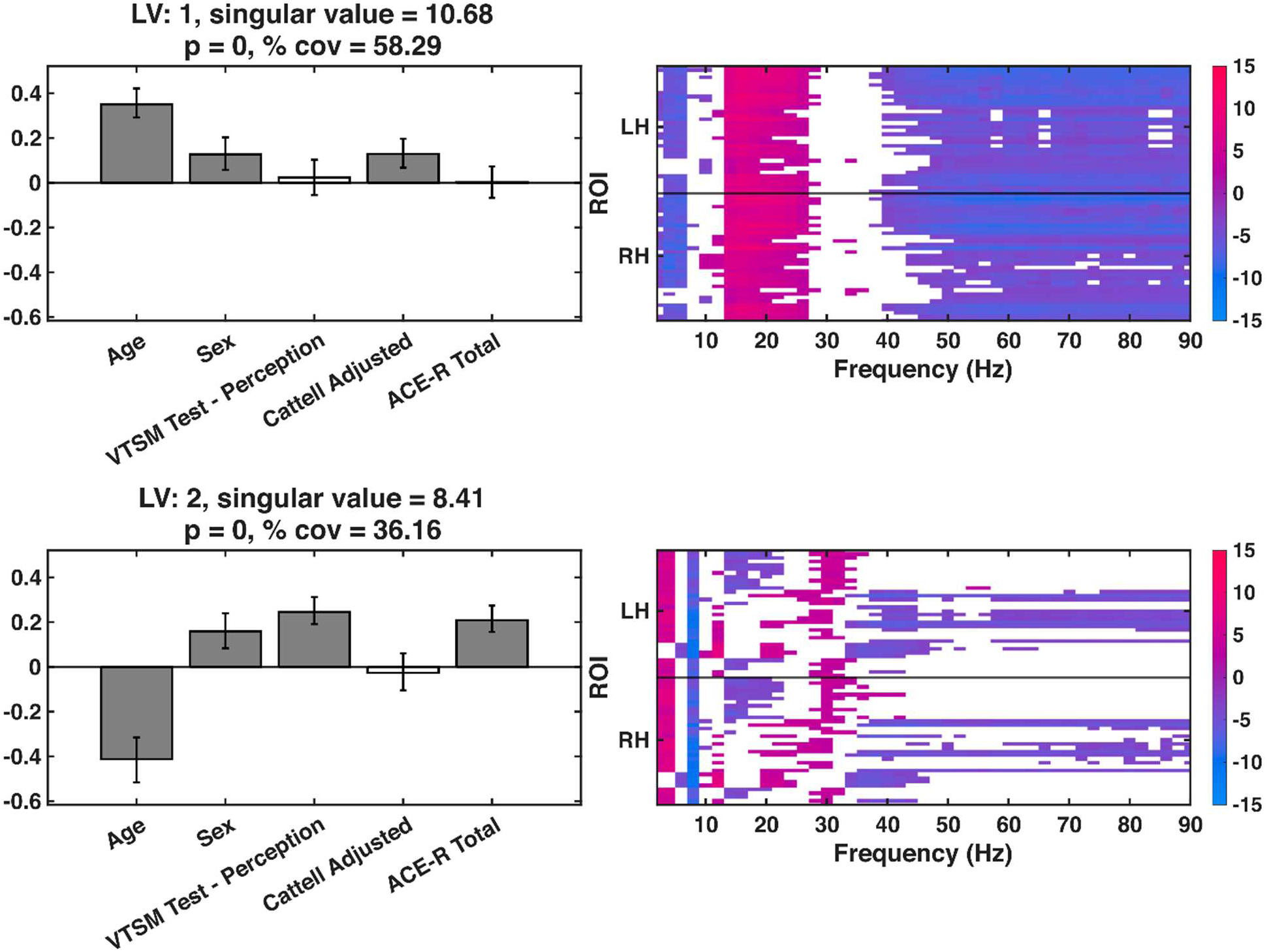
Results of the power spectrum behavioural partial least squares analysis. Each significant LV is represented by two plots: 1) a bar plot of the correlation of each behaviour to the LV ± a 95% confidence interval and 2) a color plot of the bootstrap ratios for the power spectrum from each region of interest (68 total, 34 in each hemisphere; RH = Right hemisphere, LH = Left hemisphere).

### 3.4 Age Grouped PLS

To investigate if the relationships noted in the PLS analysis were consistent across behaviours, a grouped PLS analysis was performed to allow for correlations with the LV to vary between groups. The age grouped behaviour PLS analysis separated age into 3 groups. This analysis revealed two significant LVs that account for 88.95% of the covariance (Figure 3A). Notably the first and second LVs in this model related to the first two LVs in the ungrouped PLS analysis (cos(LV1_age grouped_ X LV1 _ungrouped_) = 0.90 and cos(LV1_age grouped_ X LV2_ungrouped_) = -0.42 & cos(LV2_age grouped_ X LV1 _ungrouped_) = 0.35 and cos(LV2_age grouped_ X LV2 _ungrouped_) = 0.81), indicating that the LVs present in this analysis captured variability from both LVs in the ungrouped analysis (Figure 5A). In the first LV from the grouped PLS age was correlated in both the young and middle-aged participants, but not the old participants (Figure 5A & B). Instead, sex differences become associated with this LV in the middle-aged and old participants, whereas the sex effect for young participants loaded onto the second LV (Figures 5A).

**Figure 5:**
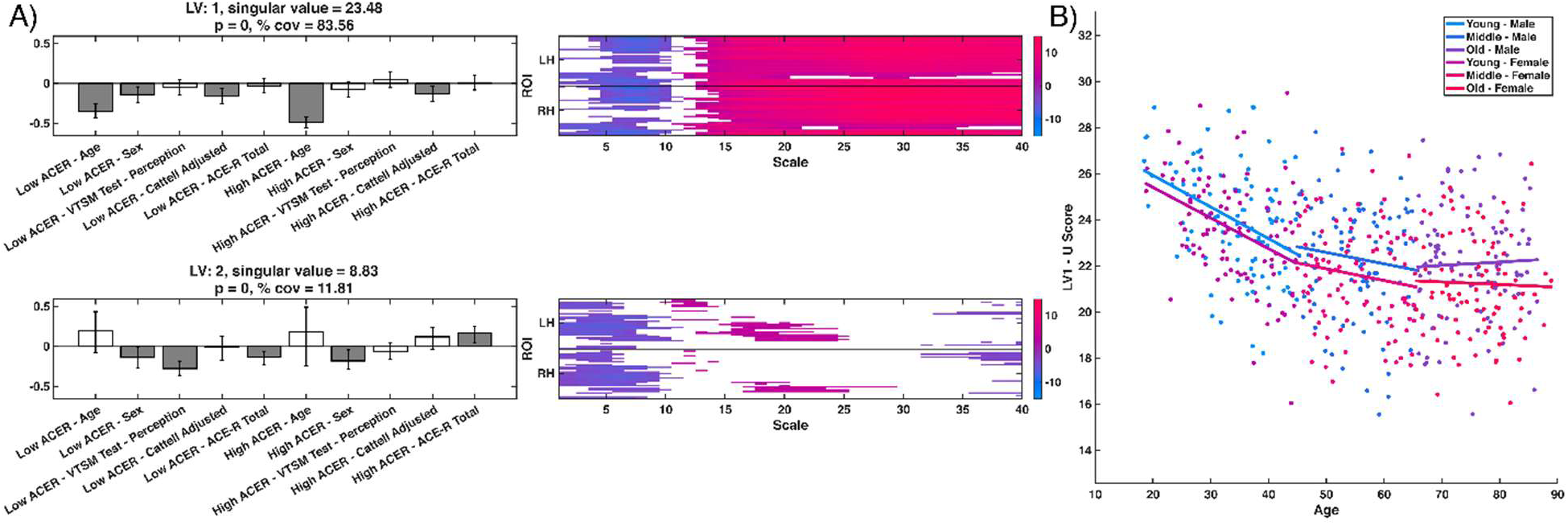
A) Results of the age grouped MSE behavioural partial least squares analysis. Each significant LV is represented by two plots: 1) a bar plot of the correlation of each behaviour to the LV ± a 95% confidence interval and 2) a color plot of the bootstrap ratios for the MSE curves from each region of interest (68 total, 34 in each hemisphere; RH = Right hemisphere, LH = Left hemisphere). Figure 3B is a scatterplot representing each participant’s score on LV1. Participants are grouped by age and sex, represented by colour. A linear regression was used to overlay a line of best fit for each group.

The age-grouped PLS analysis on the power spectrum also revealed two significant LVs that account for 86.91% of the covariance explained by the model (Figure 6A). The first and second LVs in this model covaried with both significant LVs in the ungrouped PLS analysis (cos(LV1_age grouped_ X LV1 _ungrouped_) = 0.93 and cos(LV1_age grouped_ X LV2_ungrouped_) = 0.34 & cos(LV2_age grouped_ X LV1 _ungrouped_) = -0.22 and cos(LV2_age grouped_ X LV2 _ungrouped_) = 0.74), confirming the strong age effect (Figure 6A). Similar to the MSE analysis, the first LV from the age-grouped PSD PLS analysis was correlated with age in young participants, but not the middle-aged or old participants (Figure 6A & B). Instead, sex differences become associated with this LV in the middle-aged and old participants. In contrast, the age and sex effects for young participants are split across both LV from this grouped model (Figures 6A).

**Figure 6:**
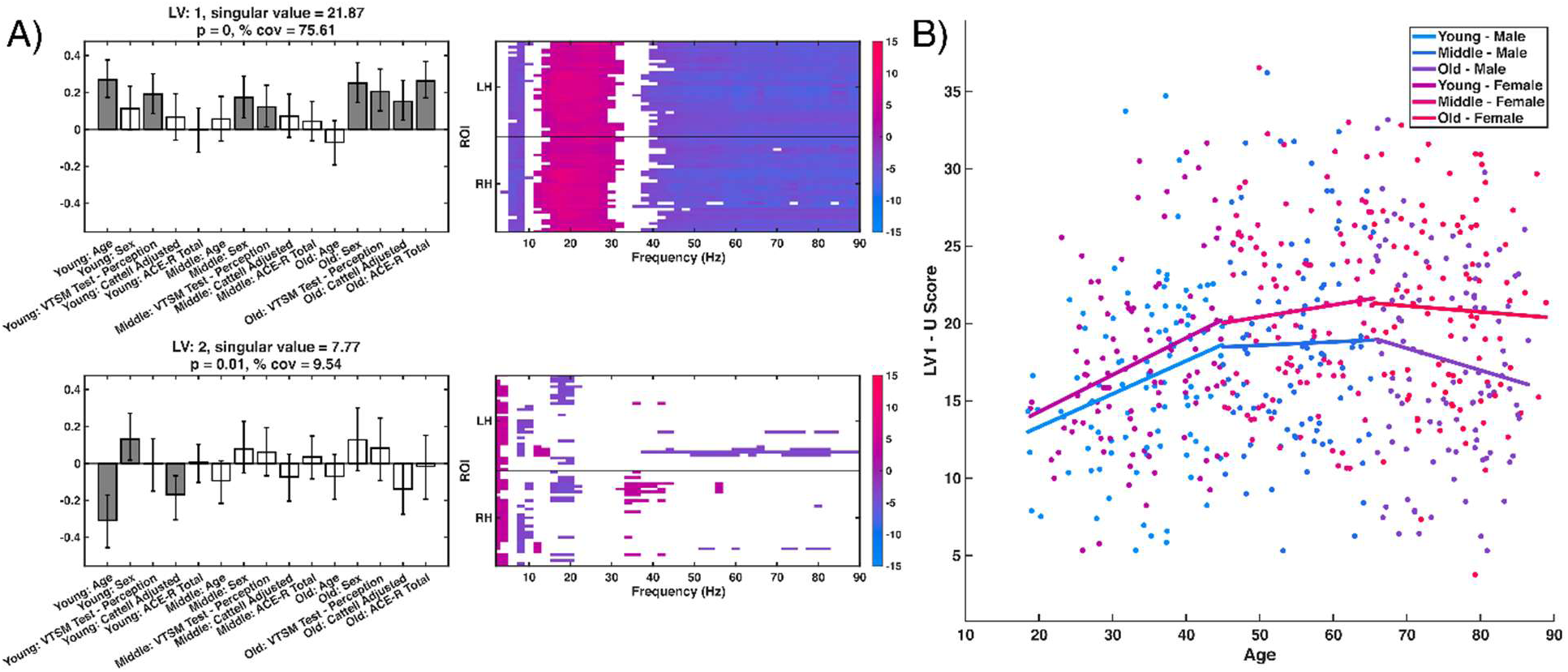
A) Results of the age grouped power spectrum behavioural partial least squares analysis. Each significant LV is represented by two plots: 1) a bar plot of the correlation of each behaviour to the LV ± a 95% confidence interval and 2) a color plot of the bootstrap ratios for the power spectrum from each region of interest (68 total, 34 in each hemisphere; RH = Right hemisphere, LH = Left hemisphere). B) A scatterplot representing each participant’s score on LV1. Participants were grouped by age and sex, represented by colour. A linear regression was used to overlay a line of best fit for each group.

### 3.5 Sex grouped PLS

When treating sex as a grouping variable in a PLS analysis of the MSE data, the test yielded one significant LV and one marginally significant LV that account for 93.76% of the covariance explained by the model (LV1: p = 0.00, LV2: p = 0.06; Supporting Information Figure 1). Notably, the first and second LVs in this model showed the highest cosine similarity with the corresponding LV in the ungrouped PLS analysis (cos(LV1_sex grouped_ X LV1 _ungrouped_) = 1 and cos(LV2 _sex grouped_ X LV2 _ungrouped_) = 0.93), confirming that the LVs described in the ungrouped test are consistent within each sex.

The sister PLS analysis on the PSD data revealed two significant LVs that account for 91.04% of the covariance explained by the model (Supporting Information Figure 2). Notably the first and second LVs in this model had high cosine similarities with both LVs in the ungrouped PLS analysis (cos(LV1_sex grouped_ X LV1 _ungrouped_) = 0.89, cos(LV1_sex grouped_ X LV2 _ungrouped_) = -0.45, cos(LV2_sex grouped_ X LV1 _ungrouped_) = 0.40 and cos(LV2 _sex grouped_ X LV2 _ungrouped_) = 0.85) suggesting that the effects captured in both LVs of the ungrouped test are expressed differently in each sex. The correlations of the behaviours with the first and second LVs in this analysis align well with the ungrouped test, whereby age and fluid intelligence correlate with LV1 and age, visual short-term memory and generalized cognitive function correlate with LV2. There only discernable differences between the sexes, is that visual short-term memory and generalized cognitive function are associated with both LVs in female participants. However, the representation of these effects in the cortex is a mix of the patterns presented in the ungrouped analysis. The cortex-wide effects in the beta and gamma bands of the ungrouped test were more regional in this analysis (Figures 4 and Supporting Information Figure 2) and the beta response seen in LV2 of the ungrouped analysis was largely absent in this analysis (Figure 4 and Supporting Information Figure 2).

### 3.6 Behaviour Grouped PLS

We tested whether the aging effects on MSE were expressed consistently across levels of fluid intelligence, visual short-term memory and generalized cognitive function based on a median split of the relevant test score. The analysis of the adjusted Cattell scores revealed two significant LVs that account for 95.67% of the covariance explained by the model which had high cosine similarity to their corresponding significant LVs in the ungrouped analysis (cos(LV1_Cattell grouped_ X LV1 _ungrouped_) = 1 and cos(LV2 _Cattell grouped_ X LV2 _ungrouped_) = 0.99. The VTSM PLS analysis revealed two significant LVs that account for 93.75% of the total variance and aligned well with the ungrouped test (cos(LV1_Cattell grouped_ X LV1 _ungrouped_) = 1 and cos(LV2 _Cattell grouped_ X LV2 _ungrouped_) = 0.93). In the ACE-R PLS analysis, both significant LVs accounted for 95.05% of the total variance and also aligned well with the ungrouped test (cos(LV1_Cattell grouped_ X LV1 _ungrouped_) = 0.99 and cos(LV2 _Cattell grouped_ X LV2 _ungrouped_) = 0.97.

Akin to the MSE analysis, a median split of the three behaviours of interest (adjusted Cattell, VTSM and ACE-R) was used to test the consistency of effects seen in the ungrouped test of the PSD data. The analysis of the adjusted Cattell scores revealed two significant LVs that account for 92.15% of the covariance explained by the model that had high cosine similarity to the respective LVs in the ungroup test analysis (cos(LV1_Cattell grouped_ X LV1 _ungrouped_) = 0.99 and cos(LV2 _Cattell grouped_ X LV2 _ungrouped_) = 0.99). The VTSM PLS analysis revealed two significant LVs that account for 88.18% of the total variance and aligned well with the ungrouped test (cos(LV1_Cattell grouped_ X LV1 _ungrouped_) = 0.99 and cos(LV2 _Cattell grouped_ X LV2 _ungrouped_) = 0.97). In the ACE-R PLS analysis, both significant LVs accounted for 90.50% of the total variance and aligned well with the ungrouped test (cos(LV1_Cattell grouped_ X LV1 _ungrouped_) = 0.98 and cos(LV2 _Cattell grouped_ X LV2 _ungrouped_) = 0.97). Collectively, the high cosine similarity between LVs of the grouped analyses and ungrouped analysis in both the MSE and PSD data suggest that the effects noted in the ungrouped PLS analyses are consistent across the cognitive behaviours tested.

### 3.7 Comparing MSE to PSD results

One of the advantages of MSE is it’s ability to characterize both the linear and non-linear dynamics in a time-series, in contrast to power spectral decompositions that are only sensitive to linear auto-correlations. The impact of non-linear effects in the MEG data was tested by running PLS on MSE curves calculated from phase-randomized time series. This analysis revealed two significant LVs that account for 97.70% of the covariance explained by the model (Supporting Information Figure 3A). Notably the first and second LVs in this model have high cosine-similarities with the corresponding LV in the original PLS analysis (cos(LV1_phase randomized_ X LV1 _ungrouped_) = 0.99 and cos(LV2 _phase randomized_ X LV2 _ungrouped_) = 0.98) suggesting that the differences observed in the initial analyses were linear. In addition, a task PLS analysis comparing the phase-randomized data to the original timeseries demonstrated that there is increased entropy in the phase-randomized signal in comparison to the unaltered data, consistent with previous literature (Supporting Information Figure 3B&C; Courtiol et al., 2016). Furthermore, all behaviour PLS analyses on the phase-randomized data produced nearly identical LVs to the original data, further reinforcing that the MSE effects reflect linear autocorrelations in the data.

Given that non-linear dynamics are not significantly contributing to the aging trends in the MSE data, the variance captured by both the MSE and PSD data should be able to explain the linear-autocorrelations related to age. However, when comparing the alignment of the behavioural LVs between the MSE and power spectrum analyses, the cosine similarities of the behavioural weights demonstrated that the age and sex effects expressed on LV1 and LV2 respectively of the MSE analysis are spread across both LVs of the power spectrum analysis (cos(LV1_power spectrum_ X LV1_MSE_) = -0.84 and cos(LV1_power spectrum_ X LV2_MSE_) = 0.54 & cos(LV2_power spectrum_ X LV1 _MSE_) = 0.54 and cos(LV2_power spectrum_ X LV2_MSE_) = 0.84). This finding suggests that the variability captured by the LVs in the power spectrum and MSE curve behavioural PLS analyses reflects similar effects in the data; however, these effects are mapped differently to the behavioural variables of interest. Specifically, the aging and sex effects are spread across both LVs in the power spectrum, whereas they are more distinctly separated within both LVs of the MSE analysis.

## 4 Discussion

This study replicated and extended previous findings on how multiscale entropy (MSE) changes across the adult lifespan. Using a large, previously characterized MEG sample, we addressed our first objective, validating the classic MSE aging profile, which includes increased fine-scale and decreased coarse-scale entropy with age. Furthermore, in accordance with our second objective, we demonstrated that these effects vary systematically by sex and, to a lesser extent, by cognitive performance. However, sex did not moderate the relationship between fluid intelligence and MSE. To address our final objective, the MSE results were compared with the power spectrum, revealing that age effects are largely driven by linear autocorrelations in the data, yet the MSE yielded unique summaries that highlighted sex-based differences. Together, these results strengthen the case for MSE as a sensitive, population-level marker of functional brain aging. There were differences in MSE curves between female and male participants, highlighting the potential for MSE as a measure that can differentiate age-related changes in network dynamics between demographic variables (Figure 3).

Interestingly, an interaction between age and sex differences appear to modulate trajectories of MSE across the adult lifespan (Figures 1B&C), potentially reflecting distinct influences on large-scale network integration. Brain derived entropy metrics have been shown to be sensitive to sex effects, whereby it has been noted that entropy is higher in women than men using a wide variety of entropy metrics applied to resting state EEG, MEG and functional MRI datasets (Ahmadi et al., 2013; Dong et al., 2018; Fernández et al., 2012; Shumbayawonda et al., 2019; Wang, 2021; Zhao et al., 2024). However, the interaction between sex and age has not been clearly elucidated as previous investigations have arrived at conflicting results concerning whether this interaction exists (Fernández et al., 2012; Sokunbi et al., 2015; Yao et al., 2013; Zhao et al., 2024). In contrast to this study, these investigations all use entropy metrics that do not consider the temporal organization of brain activity. To date, we are only aware of one investigation of sex-based differences using a multivariate version of MSE, which demonstrates that female individuals have increased fine scale entropy and decreased coarse scale entropy relative to male individuals (Lewandowska et al., 2023). This effect is consistent with the MSE results seen in the middle and old age groups where the MSE effects commonly associated with aging seem to become more related with sex (Figure 3; Lewandowska et al., 2023; McIntosh, 2019). This study builds on the results of (Lewandowska et al., 2023) by suggesting that these sex effects are more prevalent as people age and that entropy changes across the lifespan reflect a quadratic or asymptotic shape, consistent with results of Fernández et al. 2012 ^58,63^. Furthermore, the onset of these changes becomes apparent in the middle age group aligning with the crossover point (50 years old) between lifespan entropy trajectories for male and female individuals reported by (Yao et al., 2013) and, notably, the mean age at natural menopause (Schoenaker et al., 2014). These changes may therefore be reflective of decreased sex-hormones in post-menopausal female individuals, estrogen is known to have a neuroprotective effect, and the onset of these changes might be reflective of increasing dementia following menopause (Pertesi et al., 2019; Pike et al., 2009). Despite the multiple parallel findings regarding the effects of age and sex on brain-derived entropy measures, the inconsistent reporting of the age and sex interaction warrants further study.

Fluid intelligence, generalized cognitive function and short-term memory were also associated with the two dominant MSE components. Higher fluid intelligence correlated with the aging-related MSE pattern, while better short-term memory and generalized cognitive function tracked the sex-related component. In our data, sex did not moderate the relationship between MSE and fluid intelligence in contrast to the results of Dreszer et al., 2020 (Figure 2A). Such contradictory findings might be explained by differing methodological approaches used to control for aging effects on fluid intelligence. Dreszer et al., 2020 restricted participant recruitment by only considering a narrow age range, whereas we controlled for the effects of age on fluid intelligence using a linear regression. Based on our results, age remains the primary determinant of fluid intelligence even after statistical adjustment.

To complement the aging effect in MSE studied, the power spectrum was examined for common aging signatures, such as alpha slowing, decreased alpha power, and increased beta activity. These features appear but are spread across both significant latent variables (LV) in the power spectrum PLS analysis, correlating with age (Figure 3A). While bootstrap ratios across LVs seem to describe contradictory effects based on the sign of the bootstrap ratios, age negatively correlates with LV2. This negative correlation indicates that positive bootstrap ratios reflect power decreases with age. Key aging effects include cortex-wide delta power decline, decreases in theta, peak alpha slowing, increased beta, and decreased gamma power. The shift from increased beta to decreased gamma occurs near the beta-gamma boundary, with some areas showing earlier crossover (e.g., LV2), with high beta/low gamma power negatively linked to age (Figure 5). The inconsistent effect of decreasing alpha power with age contrasts against prior literature, but recent analyses of age-related changes in the power spectrum in the CamCAN dataset have also not found this expected age-related change (Gomez et al., 2013; Stier et al., 2023; Ustinin et al., 2025). Since LV2 was positively linked to sex, visual memory, and cognition, it is possible that the lack of reduced alpha power with age in this sample is being offset by the effects of other behaviours (Figure 4B).

Although these PSD changes partially overlapped with the MSE results, the mapping between domains was not one-to-one. The power spectrum and MSE curves are mathematically linked as the coarse-graining process in MSE effectively acts as a low-pass filter, so fine- (Scales 0-10) and coarse-scale (Scales 13-40) entropy effects correspond approximately to changes in frequency ranges below 90Hz (scale 0), ∼25 Hz (scale 10), ∼19Hz (scale 13) and ∼6 Hz (scale 40). MSE has the capacity to capture non-linear temporal correlations that the PSD is not sensitive to. The phase-randomization analyses further clarify the relationship between the MSE curves and the power spectrum by destroying nonlinear phase dependencies while preserving the amplitude spectrum. We found that the behaviour LVs in MSE were largely unchanged, indicating that the behaviourally relevant variance arises primarily from linear temporal correlations, which are also reflected in the power spectrum. Nevertheless, MSE differs from spectral power in how it integrates these correlations across scales: rather than partitioning variance by frequency, it quantifies the pattern irregularity of the signal over time. In this way, MSE compresses the same linear information into a compact, time-domain representation of temporal structure. Two signals can share identical spectra yet yield distinct MSE profiles, highlighting that MSE captures the organization of spectral components rather than their amplitudes, which is an essential feature of signals from complex systems (Costa et al., 2005, 2002).

These observations are consistent with the mechanistic analysis of Kosciessa et al. 2020. They demonstrated that standard MSE largely reflects spectral power at the timescales defined by coarse graining, linking entropy values directly to the spectral slope of EEG and MEG data (Kosciessa et al., 2020). Our surrogate analyses converge with this interpretation. The persistence of behavioural covariance patterns after phase randomization suggests that the behaviourally relevant variance in MSE is rooted in these spectrally determined, linearly correlated temporal structures. This relationship is further reinforced by the high degree of similarity of behavioural covariance patterns across the two metrics, MSE and PSD, in this dataset. For both metrics, the effects captured in the ungrouped analysis were consistent across the range of each behaviour of interest (fluid intelligence, generalized cognitive function and visual short-term memory). Further, the age grouped analysis highlights a trend where the pattern of brain data expressed on the first LV in both MSE and PSD data represents an aging effect in young participants that changes to largely be an expression of sex effects in old participants (Figures 3A & 6A).

Despite the high degree of alignment between MSE and PSD, prior work has also shown that the relationship between the variance in MSE and linearly correlated temporal structures of the PSD is not universal. In a study of childhood development, (McIntosh et al., 2008) demonstrated that phase randomization can alter MSE and related measures of signal dimensionality even when the power spectrum remains constant, implying sensitivity to temporal dependencies that go beyond spectral content. This apparent discrepancy could reflect differences in the underlying dynamics. This divergence suggests that while MSE and spectral power share variance related to linear autocorrelations, MSE provides a higher-order summary of temporal organization that cannot be reduced to frequency content alone. The effects of sex in this dataset exemplify these discrepancies, whereby the patterns in the variability captured in the MSE curves are consistently represented across sex, however, in the power spectrum, the representation of the behavioural effects in the cortex becomes more regional. These divergent summaries of the covariations between a demographic variable, sex, and the same brain activity highlight complementary information that MSE can provide to a frequency-based analysis. In this light, our results suggest that resting-state MEG signals, such as those in the CamCAN dataset, primarily lie within the linear regime described by (Kosciessa et al., 2020), however, this relationship may be impacted by different cognitive, demographic or developmental contexts that may invoke the more metastable dynamics captured by sex in this analysis and by the results of (McIntosh et al., 2008).

A similar perspective is presented by (Shen et al., 2021), who examined multiscale entropy and spectral power across a large, heterogeneous clinical EEG sample. Using the same PLS framework applied here, they found that MSE differentiated both neurological and non-neurological comorbidities, whereas spectral power did not (Shen et al., 2021). The divergence between the two analyses emerged despite both measures being derived from the same underlying signals, reinforcing the view that MSE and spectral metrics provide complementary but non-redundant representations of neural dynamics. In line with our findings, this suggests that even when MSE is primarily influenced by linear spectral structure, its integration of temporal dependencies across scales can reorganize that information to emphasize clinically or behaviourally relevant dimensions of variability. An example of this re-organization in the current dataset is the summarization of sex effects in the CamCAN data, which shows consistent behavioural effects on MSE within sex in contrast to the power spectral data. All told, these findings point to a continuum: from spectral-dominant regimes in resting data, through mixed regimes in clinical populations, to phase-structured, metastable dynamics in active cognition.

### 4.1 Conclusion

This MSE analysis of the resting state CamCAN data confirmed the previously characterized aging profile of increased fine-scale and decreased coarse-scale entropy. Interestingly, this profile was largely predicted by age in the younger participants, and, as participants aged, it became more strongly associated with sex. These effects were consistently present in both the MSE curves and PSD analyses, and in combination with the phase randomization analysis, reveal that these trends are driven by linearly correlated temporal structures that are equivalently captured by both analysis approaches. However, there are several contexts in which MSE’s capacity to provide a higher-order summary of temporal organization highlights unique information, as demonstrated by the divergent results of sex in this analysis or in the context of the nature of comorbidities in heterogeneous disease states. Cumulatively, these findings suggest that even while MSE and PSD reflect linear spectral structures, MSE’s sensitivity to temporal dependencies can emphasize unique relationships between neural activity and relevant clinical or behavioural contexts.

## CRediT statement

**Jack P. Solomon**: Conceptualization, Methodology, Software, Validation, Formal Analysis, Writing – Original Draft, Writing – Review and Editing, Visualization, **Simon G. J. Dobri**: Methodology, Software, Validation, Formal Analysis, Writing – Review and Editing, **Kelly Shen**: Conceptualization, Resources, Data Curation, Project Administration, **Vasily A. Vakorin**: Methodology, Validation, Resources, Data Curation, Writing – Review and Editing, **Sylvain Moreno**: Resources, Data Curation, Writing – Review and Editing, Supervision, Project Administration, Funding acquisition, **Anthony R. McIntosh**: Conceptualization, Methodology, Validation, Resources, Data Curation, Visualization, Supervision, Project Administration, Funding acquisition, Writing – Review and Editing

## Acknowledgements

This research was enabled in part by support provided by the Research Computing Group at Simon Fraser University and computational resources provided by Digital Research Alliance of Canada.

## Data availability statement

The CamCAN dataset used in the analysis is available at (https://camcan-archive.mrc-cbu.cam.ac.uk/dataaccess/) and the MRI-MEG co-registration files for the CamCAN dataset are available at (https://github.com/tbardouille/camcan_coreg). The code used for this analysis is available on the McIntosh-Lab GitHub organization which includes the MEG processing pipeline (https://github.com/McIntosh-Lab/Solomon_2026_MEG_pipeline) and subsequent MSE analysis (https://github.com/McIntosh-Lab/camcan_mse_analysis).

## Funding

Funding for this project was provided by Natural Sciences and Engineering Research Council of Canada Discovery Grant *RGPIN-2018-04457* and the Canadian Institutes of Health Research Project Grant *PJT-168980*, awarded to Dr. Randy McIntosh. Additional funding was provided for Dr. Jack Solomon as Michael Smith Health Research BC / Alzheimer’s BC Research Trainee.

## Disclosure Statement

No conflicts of interest to report

## Notes

https://github.com/McIntosh-Lab/Solomon_2026_MEG_pipeline

https://github.com/McIntosh-Lab/camcan_mse_analysis

### Competing Interest Statement

The authors have declared no competing interest.

