## Supplementary figures and images for "Interactions between age and sex in multiscale entropy and spectral power changes across the lifespan"

### Supporting Information Figure 1

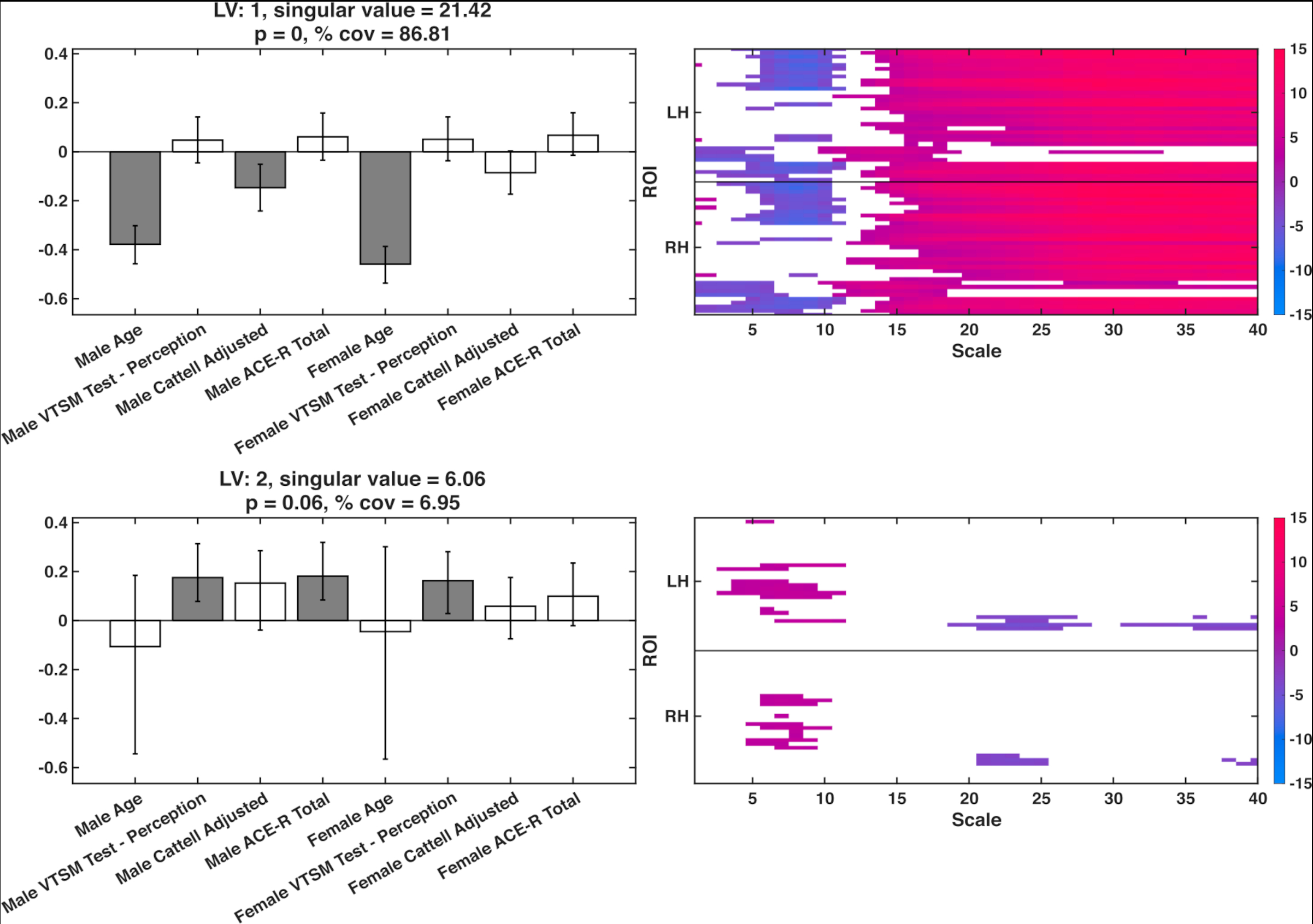

### Supporting Information Figure 2

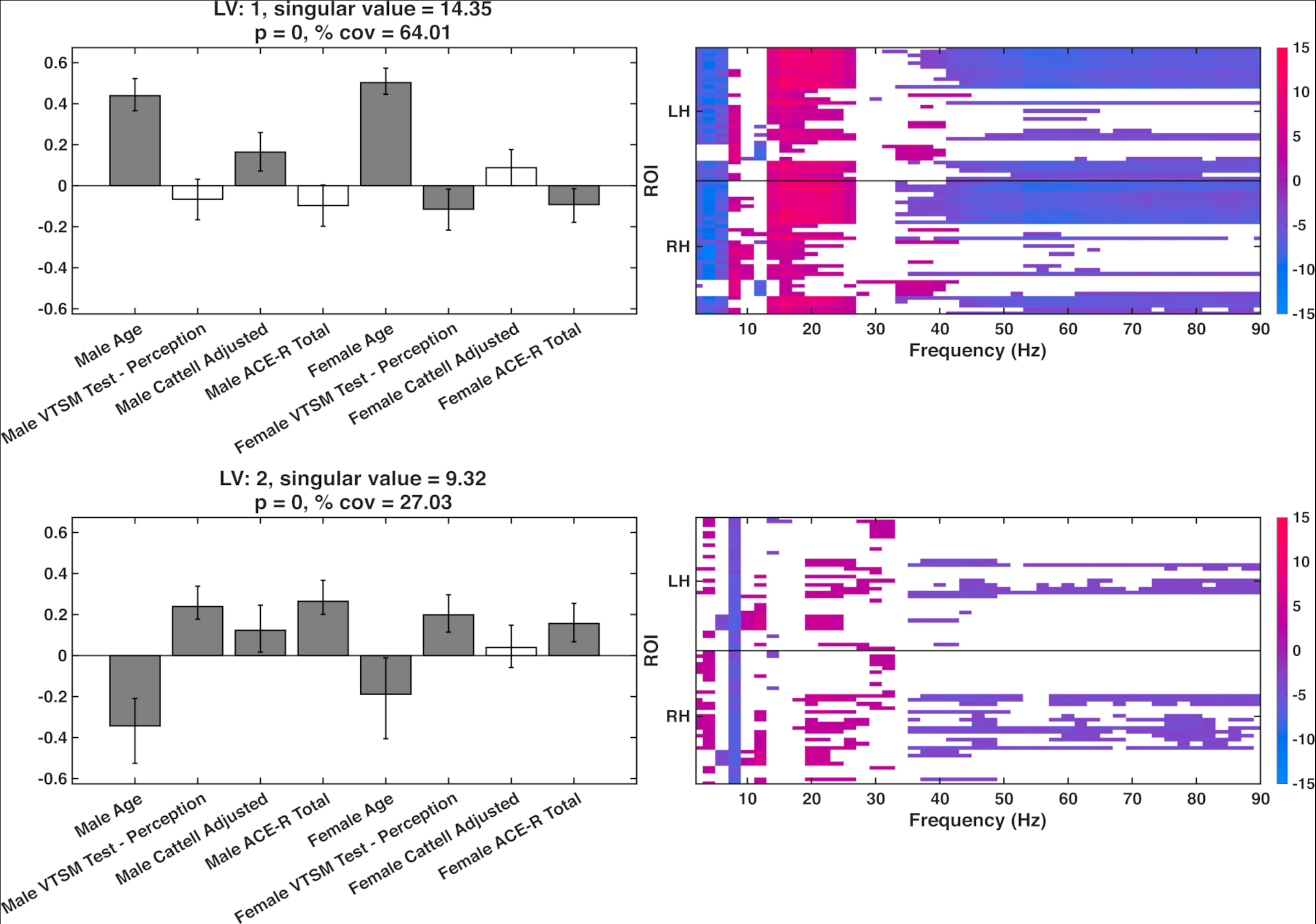

### Supporting Information Figure 3

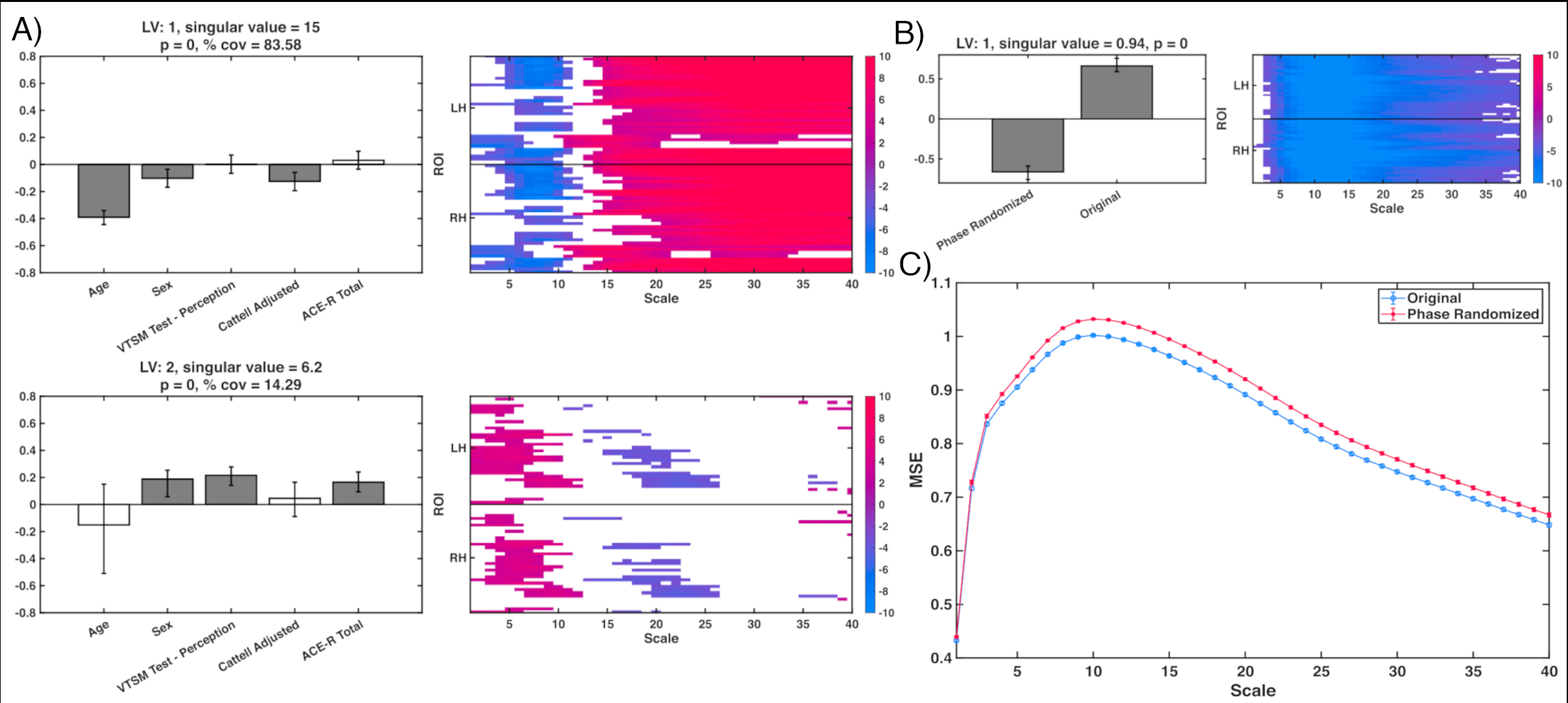
